# PRMT5 regulates ovarian follicle development by facilitating *Wt1* translation

**DOI:** 10.1101/2021.04.04.438394

**Authors:** Min Chen, Fangfang Dong, Min Chen, Zhiming Shen, Haowei Wu, Changhuo Cen, Xiuhong Cui, Shilai Bao, Fei Gao

## Abstract

Protein arginine methyltransferase 5 (*Prmt5*) is the major type II enzyme responsible for symmetric dimethylation of arginine. Here, we found PRMT5 was expressed at high level in ovarian granulosa cells of growing follicles. Inactivation of *Prmt5* in granulosa cells resulted in aberrant follicle development and female infertility. In *Prmt5-*knockout mice, follicle development was arrested with disorganized granulosa cells in which WT1 expression was dramatically reduced and the expression of steroidogenesis-related genes was significantly increased. The premature differentiated granulosa cells were detached from oocytes and follicle structure was disrupted. Mechanism studies revealed that *Wt1* expression was regulated by PRMT5 at the protein level. PRMT5 facilitated IRES-dependent translation of *Wt1* mRNA by methylating HnRNPA1. Moreover, the upregulation of steroidogenic genes in *Prmt5*-deficient granulosa cells was repressed by *Wt1* overexpression. These results demonstrate PRMT5 participates in granulosa cell lineage maintenance by inducing *Wt1* expression. Our study uncovers a new role of post-translational arginine methylation in granulosa cell differentiation and follicle development.

## Introduction

Follicles are the basic functional units in the ovaries. Each follicle consists of an oocyte, the surrounding granulosa cells and theca cells in the mesenchyme. The interaction between oocytes and somatic cells is crucial for follicle development. Follicle maturation experiences primordial, primary, secondary, and antral follicular stages. Primordial follicles are formed shortly after birth via breakdown of oocyte syncytia. Each primordial follicle is composed of an oocyte surrounded by a single layer of flattened pregranulosa cells that remains in a dormant phase until being recruited into the primary stage under the influence of two main signaling pathways^1^. Once activated, flattened granulosa cells become cuboidal, and follicles continue to grow through proliferation of granulosa cells and enlargement of oocytes. Development of high-quality oocytes is important for female reproductive health and fertility^1–4^. Although gonadotropin, follicle-stimulating hormone (FSH) and luteinizing hormone (LH) are important for the growth of antral follicles, the early stages of follicle development are driven by a local oocyte-granulosa cell dialog. Abnormalities in this process may lead to follicle growth arrest or atresia^2, 5^.

Granulosa cells are derived from progenitors of the coelomic epithelium that direct sexual differentiation at the embryonic stage and support oocyte development postnatally^4, 6^. Theca-interstitial cell differentiation occurs postnatally along with the formation of secondary follicles. The steroid hormone produced by theca-interstitial cells plays important roles in follicle development and maintenance of secondary sexual characteristics^4^. The Wilms’ tumor (WT) suppressor gene *Wt1* is a nuclear transcription factor indispensable for normal development of several tissues. In gonads, *Wt1* is mainly expressed in ovarian granulosa cells and testicular Sertoli cells. During follicle development, *Wt1* is expressed at high levels in granulosa cells of primordial, primary and secondary follicles, but its expression is decreased in antral follicles^7^. Our previous studies demonstrated that *Wt1* is required for the lineage specification and maintenance of Sertoli and granulosa cells^8, 9^. However, the underlying mechanism that regulates the expression of *Wt1* in granulosa cells is unknown.

Protein arginine methyltransferase 5 (PRMT5) is a member of the PRMT family that catalyzes the transfer of methyl groups from S-adenosylmethionine to a variety of substrates and is involved in many cellular processes, such as cell growth, differentiation and development^10–12^. PRMT5 is the predominant type II methyltransferase that catalyzes the formation of most symmetric dimethylarginines (SDMAs) in the cells and regulates gene expression at the transcriptional and posttranscriptional levels^10^. PRMT5 forms a complex with its substrate-binding partner, the WD-repeat protein MEP50 (or WDR77), which greatly enhances the methyltransferase activity of PRMT5 by increasing its affinity for protein substrates^11^. In gonad development, inactivation of *Prmt5* specifically in primordial germ cells (PGCs) causes massive loss of PGCs^13–15^. PRMT5 promotes PGC survival by regulating RNA splicing^13^ and suppressing transposable elements at the time of global DNA demethylation^14^. In this study, we found that PRMT5 is expressed at high level in ovarian granulosa cells of growing follicles and the expression level changes with follicle development, suggesting that PRMT5 in granulosa cells plays a role in follicle development. To test the function of PRMT5 in granulosa cells, we specifically inactivated *Prmt5* in granulosa cells using *Sf1-cre*. We found that *Prmt5^flox/flox^;Sf1-cre* female mice were infertile and that follicles were arrested at the preantral stage. The expression of WT1 was dramatically reduced, and the granulosa cells in secondary follicles began to express steroidogenic genes. Further studies revealed that PRMT5 regulates follicle development by facilitating *Wt1* translation.

## Results

### Deletion of Prmt5 in granulosa cells caused aberrant ovary development and female infertility

The expression of PRMT5 in ovarian granulosa cells was examined by immunofluorescence. As shown in Fig. S1, PRMT5 (red) was expressed in oocytes, but no PRMT5 signal was detected in the granulosa cells of primordial follicles (A, A’, white arrows). PRMT5 started to be expressed in granulosa cells of primary follicles (B, B’, white arrows) and was continuously expressed in granulosa cells of secondary follicles (C, C’, white arrows), preantral follicles (D, D’, white arrows), and antral follicles (E, E’, white arrows), but its expression decreased significantly in the corpus luteum (F, F’, white arrows). To test the functions of PRMT5 in granulosa cell development, we specifically deleted *Prmt5* in granulosa cells by crossing *Prmt5^flox/flox^* mice with *Sf1-cre* transgenic mice. In *Prmt5^flox/flox^;Sf1-cre* female mice, PRMT5 expression was completely absent from granulosa cells (Fig. S2, arrows in B, D), whereas the expression of PRMT5 in oocytes and interstitial cells was not affected, suggesting that *Prmt5* was specifically deleted in granulosa cells.

**Figure S1.**
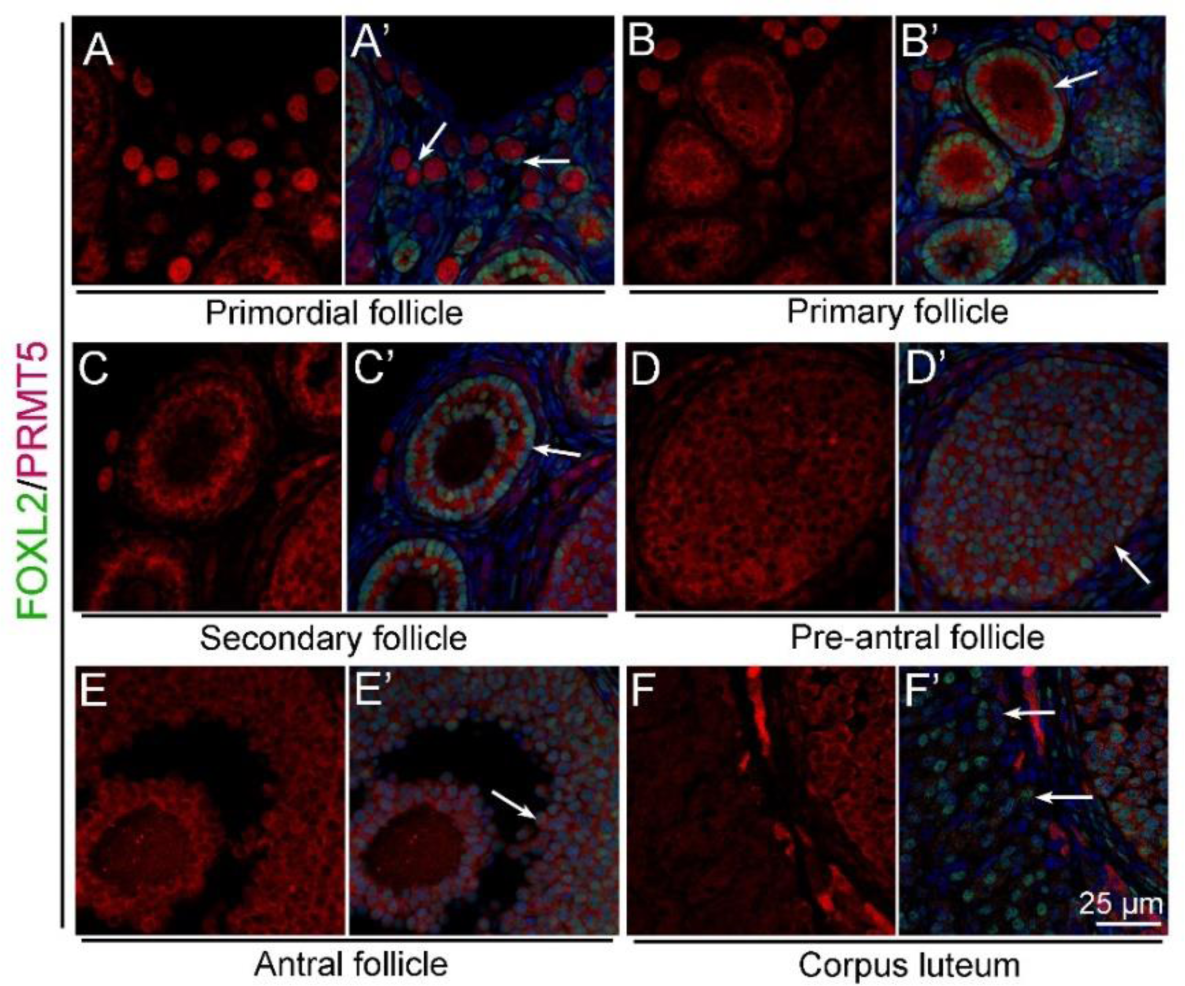
PRMT5 was expressed in granulosa cells of growing follicles. The expression of PRMT5 was examined by immunofluorescence (red), and granulosa cells were labeled with FOXL2 (green). PRMT5 was not expressed in granulosa cells of primordial follicles (A, A’, white arrows). PRMT5 was expressed in granulosa cells of primary follicles (B, B’, white arrows), secondary follicles (C, C’, white arrows), preantral follicles (D, D’, white arrows), and antral follicles (E, E’, white arrows). No PRMT5 signal was detected in the corpus luteum (F, F’, white arrows). DAPI (blue) was used to stain the nuclei.

**Figure S2.**
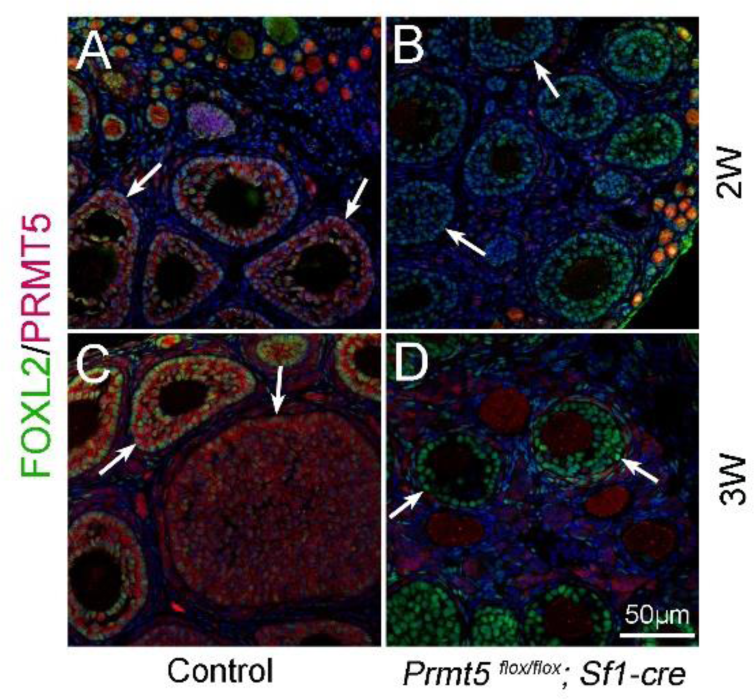
*Prmt5* was deleted in granulosa cells of *Prmt5^flox/flox^;Sf1-cre* mice. The expression of PRMT5 was examined by immunofluorescence (red), and granulosa cells were labeled with FOXL2 (green). PRMT5 protein was detected in granulosa cells of control ovaries at 2 weeks (A, white arrows) and 3 weeks (C, white arrows) after birth. No PRMT5 signal was detected in granulosa cells of *Prmt5^flox/flox^;Sf1-cre* ovaries at 2 weeks (B, white arrows) and 3 weeks (D, white arrows). DAPI (blue) was used to stain the nuclei.

No obvious developmental abnormalities were observed in adult *Prmt5^flox/flox^;Sf1-cre* mice (Fig. 1A). However, the female mice were infertile with atrophic ovaries (Fig. 1B). The results of immunohistochemistry showed growing follicles at different stages in control ovaries at 2 months of age (Fig. 1C). In contrast, only a small number of follicles and few corpora lutea were observed in *Prmt5^flox/flox^;Sf1-cre* mice (Fig. 1D). We further examined follicle development in *Prmt5^flox/flox^;Sf1-cre* mice at different developmental stages. As shown in Fig. 1, a large number of growing follicles at the primary and secondary stages were observed in *Prmt5^flox/flox^;Sf1-cre* mice (F) at 2 weeks, which was comparable to the situation in control ovaries (E). Most of the follicles were at the preantral and antral follicle stages in control mice at 3 weeks (G), whereas the development of follicles in *Prmt5^flox/flox^;Sf1-cre* mice was arrested, and aberrant granulosa cells were observed (H). The defects in follicle development were more obvious at 4 (J) and 5 weeks (L). The number of granulosa cells around oocytes was dramatically reduced, and follicle structure was disrupted.

**Figure 1.**
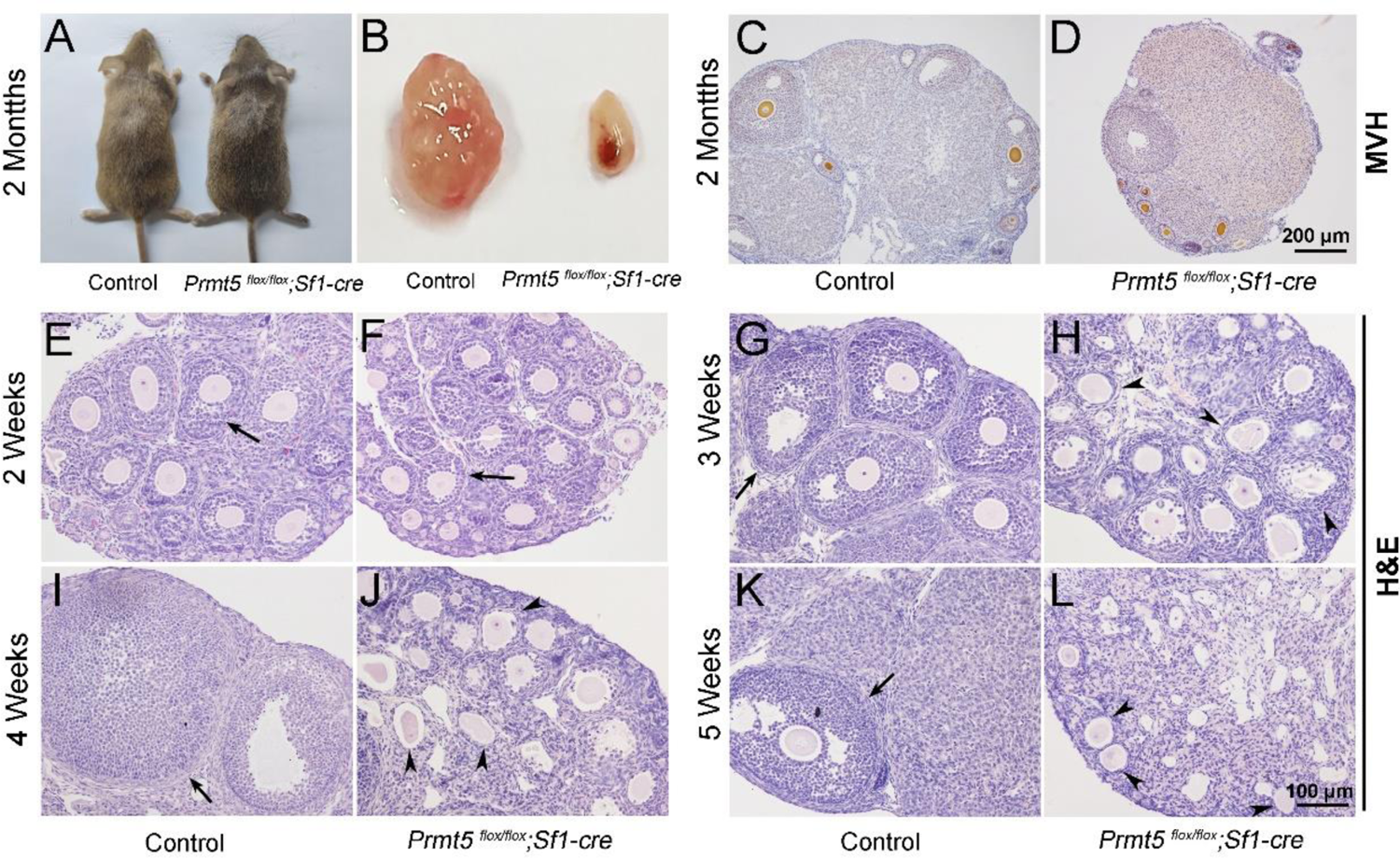
Loss of *Prmt5* in granulosa cells caused aberrant follicle development and female infertility. No developmental abnormalities were observed in *Prmt5^flox/flox^;Sf1-cre* mice (A) at 2 months of age, and the ovary size was dramatically reduced (B). Morphology of ovaries from control (C) and *Prmt5^flox/flox^;Sf1-cre* mice (D) at 2 months of age. The morphology of ovarian follicles was grossly normal in *Prmt5^flox/flox^* mice at 2 weeks (F, black arrows). Defects in follicle development were observed in *Prmt5*-mutant mice at 3 weeks (H, black arrowheads). Aberrant ovarian follicles with disorganized granulosa cells were observed in *Prmt5^flox/flox^;Sf1-cre* mice at 4 (J, black arrowheads) and 5 (L, black arrowheads) weeks of age.

### The identity of granulosa cells in Prmt5^flox/flox^;Sf1-cre mice was changed

To explore the underlying mechanism that caused the defects in follicle development in *Prmt5^flox/flox^;Sf1-cre* mice, the expression of granulosa cell-specific genes was analyzed by immunohistochemistry. As shown in Fig. 2, FOXL2 protein was expressed in the granulosa cells of both control (A, C) and *Prmt5^flox/flox^;Sf1-cre* mice (B, D) at P14 and P18. WT1 protein was expressed in granulosa cells of primary, secondary and preantral follicles in control mice at P14 and P18 (E, E’, G, G’, arrows). WT1 was also detected in the follicles of *Prmt5^flox/flox^;Sf1-cre* mice at P14 (F, arrow).

**Figure 2.**
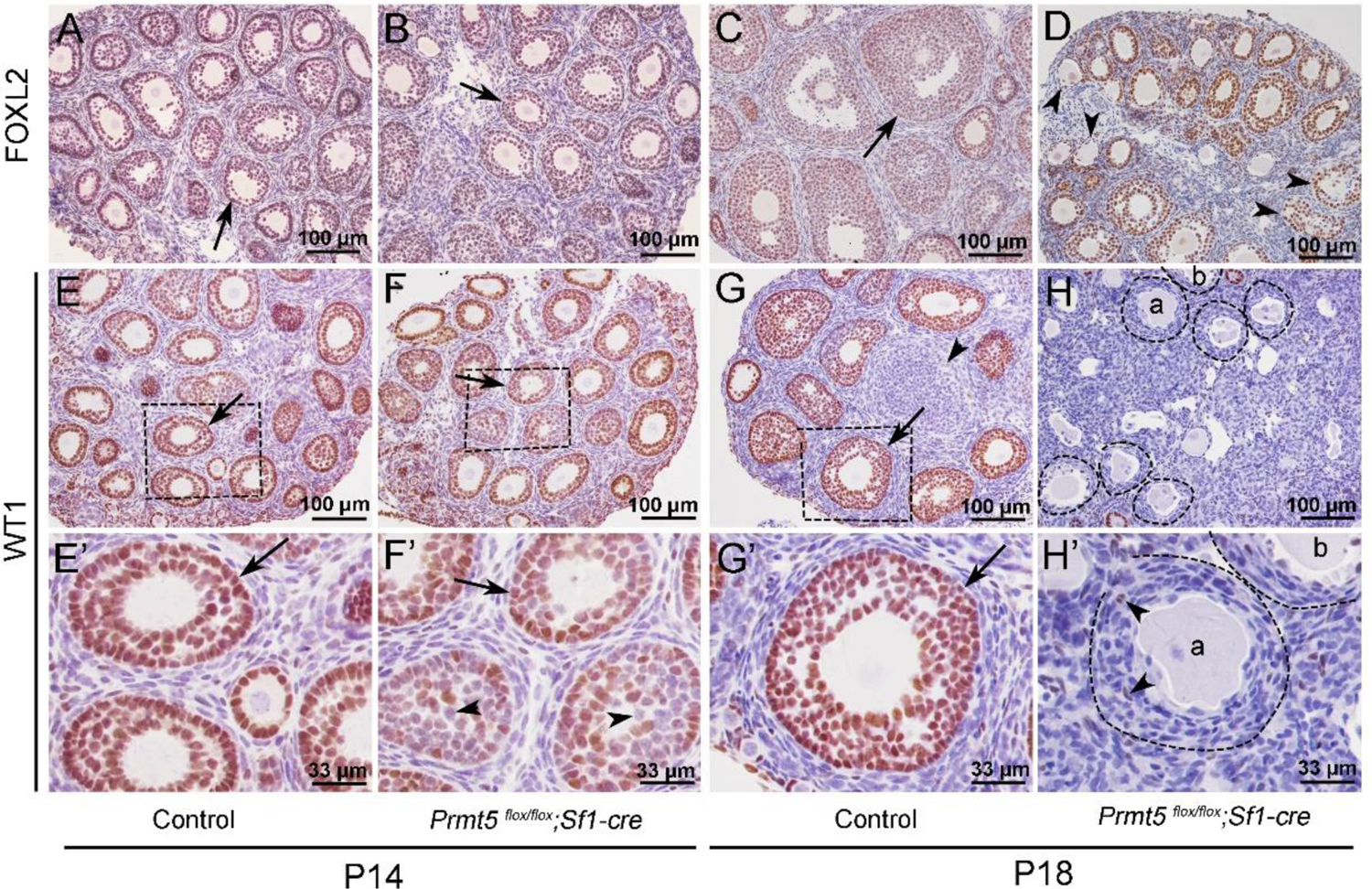
The expression of WT1 was dramatically reduced in the granulosa cells of *Prmt5^flox/flox^;Sf1-cre* mice at P18. The expression of FOXL2 and WT1 in granulosa cells of control and *Prmt5^flox/flox^;Sf1-cre* mice was examined by immunohistochemistry. FOXL2 protein was expressed in the granulosa cells of both control (A, C) and *Prmt5^flox/flox^;Sf1-cre* mice (B, D) at P14 and P18. WT1 protein was expressed in granulosa cells of primary, secondary and preantral follicles in control mice at P14 and P18 (E, E’, G, G’, black arrows). WT1 expression was absent from most granulosa cells in *Prmt5^flox/flox^;Sf1-cre* mice at P18 (H, H’); only very few granulosa cells were WT1-positive (H’, black arrowheads). E’-H’ are the magnified views of E-H, respectively.

However, not all the granulosa cells were WT1-positive; some of them were WT1-negative (F’, arrowheads). The WT1 signal was almost completely absent from the majority of granulosa cells in *Prmt5^flox/flox^;Sf1-cre* mice at P18 (H, H’); very few granulosa cells were WT1-positive (H’, arrowheads). We also found that the granulosa cells in control ovaries were cuboidal and well-organized (G’, arrow). In contrast, the granulosa cells in *Prmt5^flox/flox^;Sf1-cre* mice were flattened (H’, dashed line circle) and were indistinguishable from surrounding stromal cells. The decreased WT1 protein expression in *Prmt5*-deficient granulosa cells was also confirmed by FOXL2/WT1 double staining (Fig. S3).

**Figure S3.**
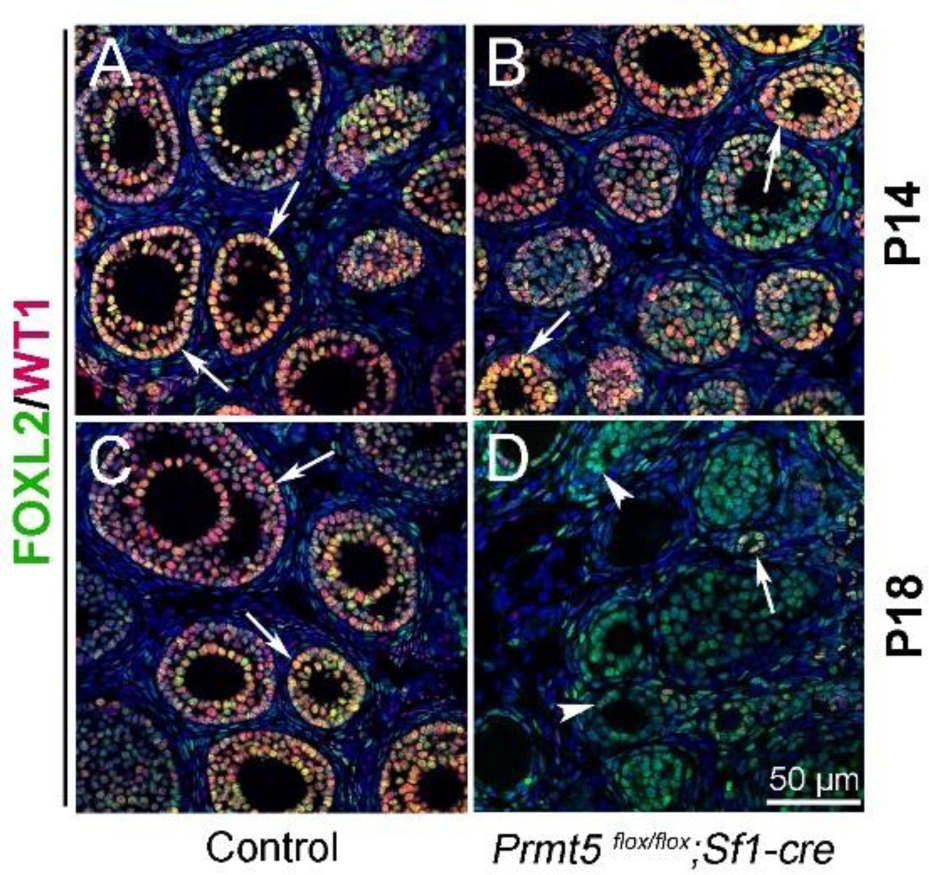
WT1 expression was decreased significantly in *Prmt5^flox/flox^;Sf1-cre* granulosa cells at P18. The expression of FOXL2 (green) and WT1 (red) in ovaries of control and *Prmt5^flox/flox^;Sf1-cre* mice at P14 and P18 was examined by immunofluorescence. WT1 expression was decreased dramatically in *Prmt5^flox/flox^;Sf1-cre* granulosa cells at P18 (D, arrowheads). Few WT1-positive granulosa cells remained (D, arrows). DAPI (blue) was used to stain the nuclei.

*Wt1* plays a critical role in granulosa cell development, and mutation of *Wt1* leads to pregranulosa cell-to-steroidogenic cell transformation^8, 9^. Therefore, we further examined the expression of steroidogenic genes in *Prmt5*-deficient granulosa cells at P18. As shown in Fig. 3, in control ovaries, 3β-HSD (3β-hydroxysteroid dehydrogenase, also known as Hsd3B1) and CYP11A1 (cytochrome P450, family 11, subfamily a, polypeptide 1, also known as P450scc) were expressed in theca-interstitial cells (A, C, arrowheads). In addition to theca-interstitial cells, 3β-HSD (B, green, arrows) and CYP11A1 (D, red, arrows) were also detected in the granulosa cells of *Prmt5^flox/flox^;Sf1-cre* mice. We also examined the expression of SF1 (steroidogenic factor 1, also known as NR5A1), which is a key regulator of steroid hormone biosynthesis^16^. As expected, SF1 was expressed only in theca-interstitial cells of control ovaries (E, red, arrowheads), whereas a high level of SF1 expression was detected in *Prmt5*-deficient granulosa cells (F, red, arrows), suggesting the identity of granulosa cells was changed. The follicle structure was destroyed as indicated by disorganized Laminin staining (H, arrows). The proliferation and apoptosis of *Prmt5*-deficient granulosa cells were analyzed by BrdU incorporation and TUNEL assays. As shown in Fig. S4, the numbers of TUNEL-positive cells and BrdU-positive granulosa cells were not changed in *Prmt5^flox/flox^;Sf1-cre* ovaries compared to control ovaries at P14 and P18.

**Figure 3.**
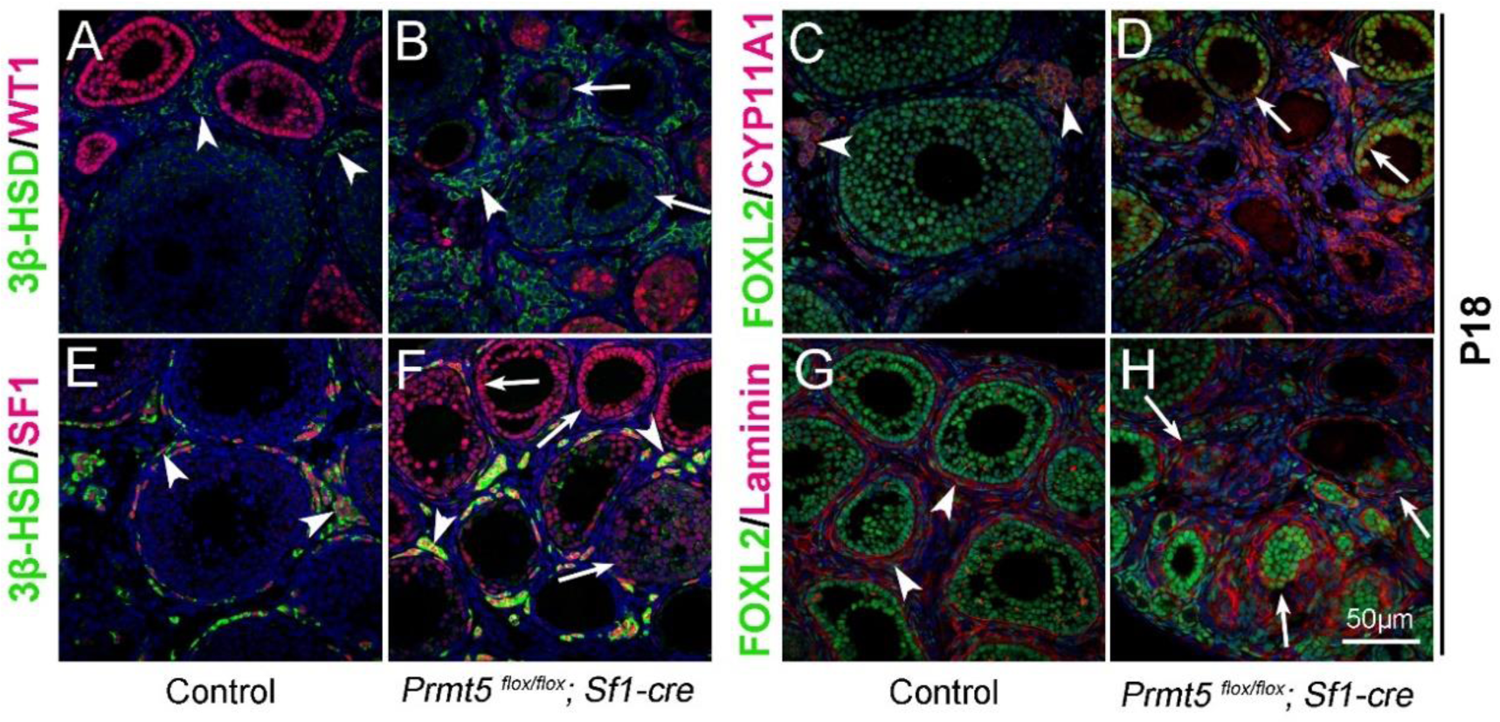
The identity of granulosa cells in *Prmt5^flox/flox^;Sf1-cre* mice was changed. The expression of 3β-HSD, WT1, FOXL2, CYP11A1, and SF1 in ovaries of control and *Prmt5^flox/flox^;Sf1-cre* mice at P18 was examined by immunofluorescence. In control ovaries, 3β-HSD (A), CYP11A1 (C), and SF1 (E) were expressed only in theca-interstitial cells (white arrowheads). In the ovaries of *Prmt5^flox/flox^;Sf1-cre* mice, 3β-HSD (B), CYP11A1 (D), and SF1 (F) were also detected in granulosa cells (white arrows). The arrows in H point to the disordered follicle structure as shown by Laminin expression. DAPI (blue) was used to stain the nuclei.

**Figure S4.**
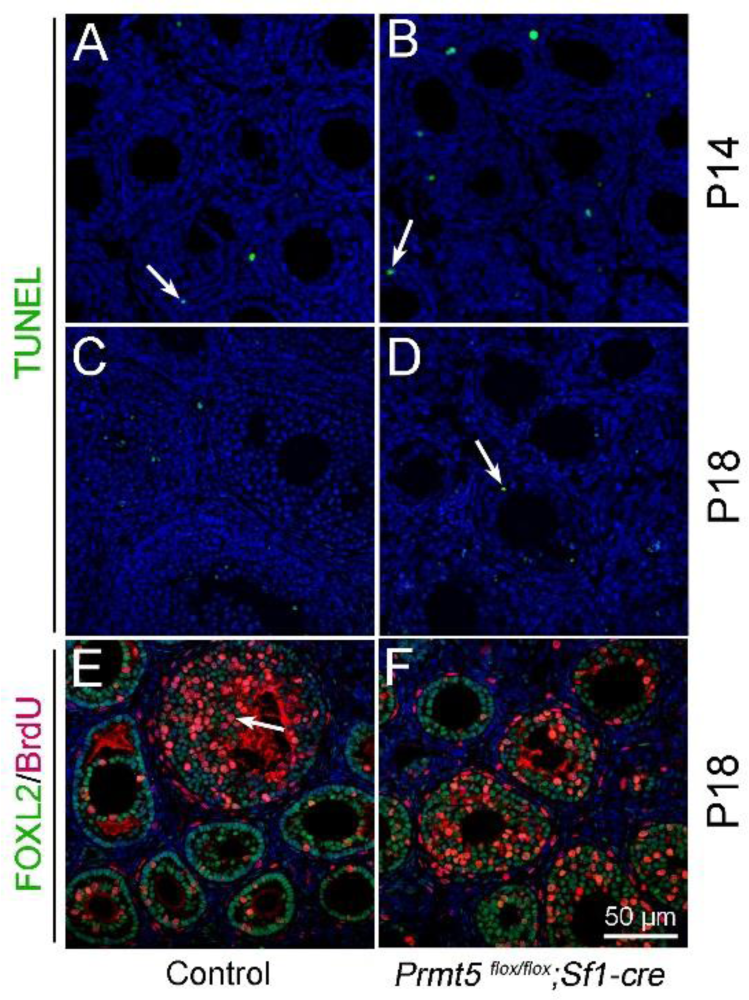
The apoptosis and proliferation of granulosa cells was not changed in *Prmt5^flox/flox^;Sf1-cre* mice at P14 and P18. Apoptosis and cell proliferation were assessed by TUNEL assay (A-D) and BrdU incorporation assay (E-F), respectively. The numbers of TUNEL-positive cells and BrdU-positive cells were not different in *Prmt5^flox/flox^;Sf1-cre* ovaries compared to control ovaries. DAPI (blue) was used to stain the nuclei.

To further confirm the above results, follicles were dissected from the ovaries of 2-week-old mice and cultured in vitro. As shown in Fig. 4, the morphology of follicles from *Prmt5^flox/flox^;Sf1-cre* mice was comparable to that of control follicles at D2. Proliferation of granulosa cells in control follicles was observed at D4, and the follicles developed to the preovulatory stage with multiple layers of granulosa cells after 9 days of culture (A-C, G-H). The granulosa cells were detached from oocytes in *Prmt5^flox/flox^;Sf1-cre* follicles at D4 (E, and a magnified view in L), and no colonized granulosa cells were observed after 9 days of culture (D-F, J-K). Most of the granulosa cells were attached to the culture dishes just like the interstitial cells, and denuded oocytes were observed after 3 days of culture (E-F, J-K, and a magnified view in L).

**Figure 4.**
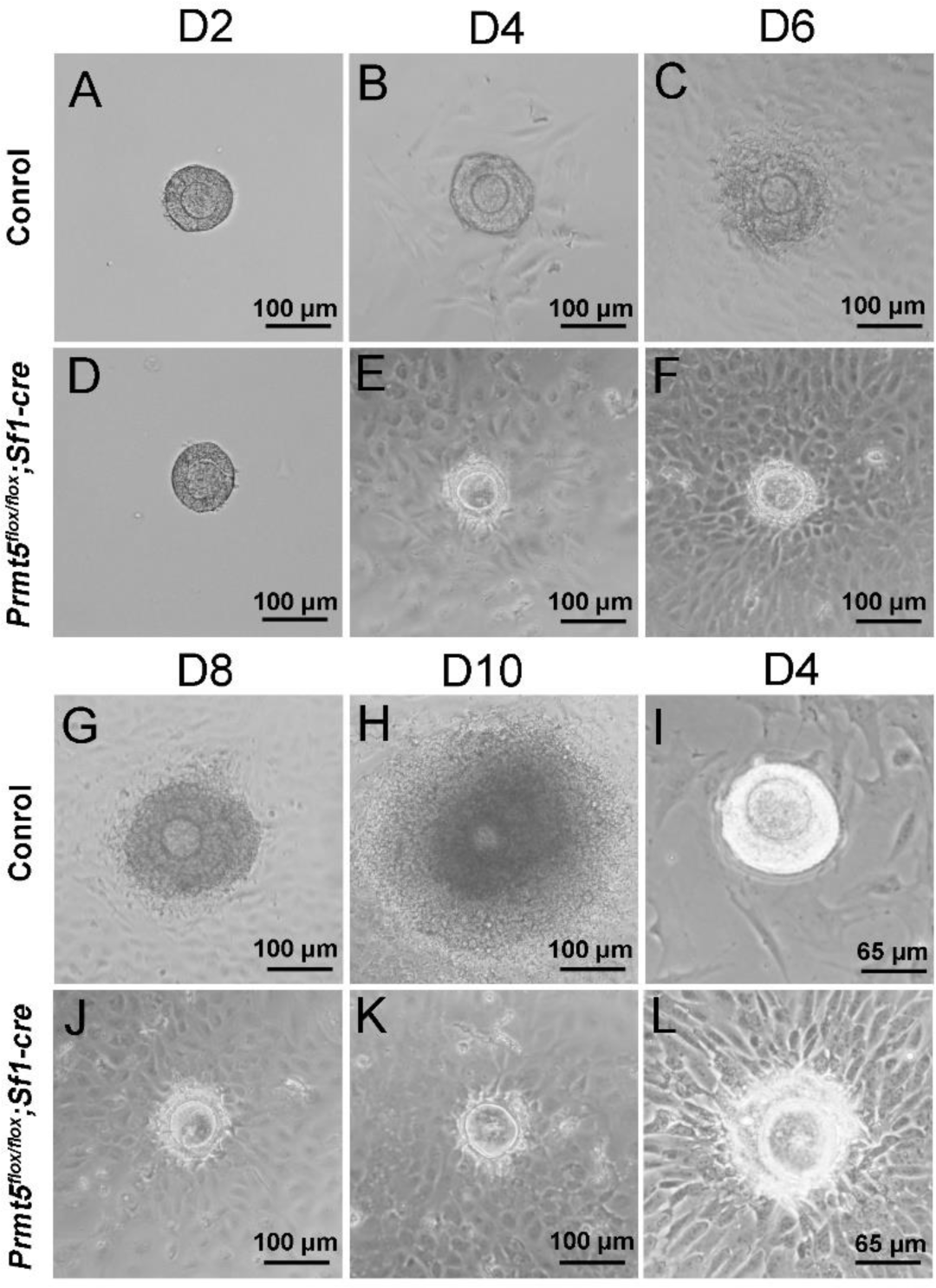
Aberrant development of in vitro-cultured *Prmt5^flox/flox^;Sf1-cre* follicles. Follicles with 2-3 layers of granulosa cells isolated from control and *Prmt5^flox/flox^;Sf1-cre* mice were cultured in vitro. After 9 days of culture, control follicles grew significantly and developed to the preovulatory stage (A-C, G-H). No obvious layers of granulosa cells were observed around oocytes (D-F, J-K), and the granulosa cells extended away from the oocytes and adhered to the dish (L). I and L are two magnified views of cultured follicles at day 4.

To further verify the differential expression of granulosa cell-specific and steroidogenic genes in *Prmt5*-deficient granulosa cells, granulosa cells were isolated at P18, and gene expression was analyzed by Western blot and real-time PCR analyses. As shown in Fig. 5 (A, B), the protein levels of PRMT5 and its interacting partner MEP50 were decreased dramatically in *Prmt5^flox/flox^;Sf1-cre* granulosa cells, as expected. The protein level of WT1 was significantly reduced in *Prmt5*-deficient granulosa cells. Surprisingly, the mRNA level of *Wt1* was not changed in *Prmt5*-deficient granulosa cells (C). FOXL2 expression was also decreased, but the difference was not significant. The expression of the steroidogenic genes CYP11A1, StAR and NR5A1 was significantly increased in *Prmt5*-deficient granulosa cells, consistent with the immunostaining results. Their mRNA levels were also significantly increased (C). We also examined the functions of PRMT5 by treating granulosa cells with the PRMT5-specific inhibitor EPZ015666. The protein level of WT1 was significantly reduced after EZP015666 treatment, whereas the mRNA level was not changed. The expression of steroidogenic genes was significantly increased at both the protein and mRNA levels after EZP015666 treatment (Fig. 5D, E, F). These results were consistent with those in *Prmt5*-deficient granulosa cells, indicating the effect of PRMT5 on granulosa cells was dependent on its methyltransferase activity. These results suggest that PRMT5 is required for maintenance of granulosa cell identity and that inactivation of this gene causes granulosa cell-to-steroidogenic cell transformation.

**Figure 5.**
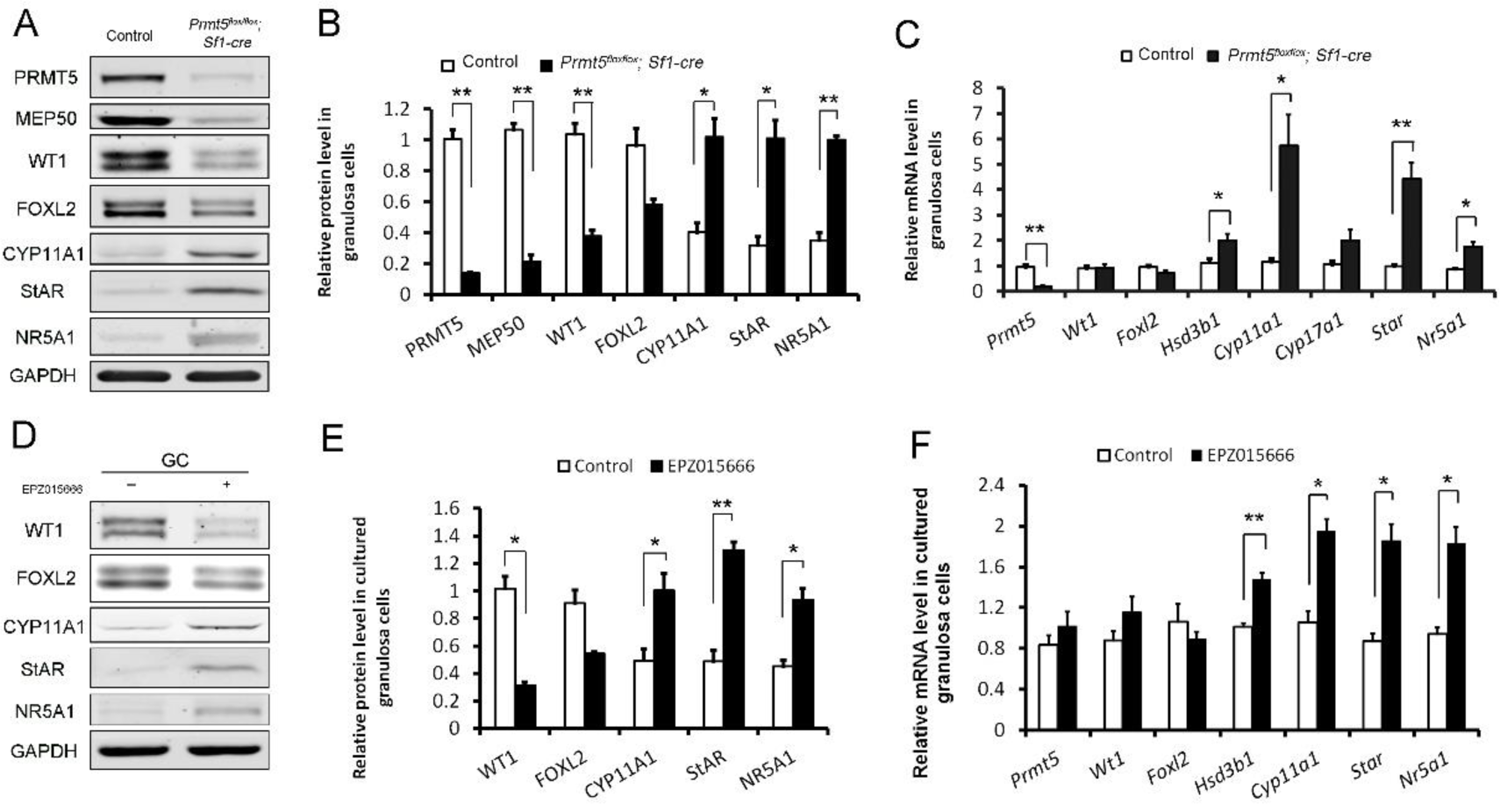
Differentially expressed genes in *Prmt5-*deficient granulosa cells. Western blot (A and B) and real-time PCR analyses (C) of the indicated genes in granulosa cells isolated from control or *Prmt5^flox/flox^;Sf1-cre* ovaries at P18. Western blot (D and E) and real-time PCR analyses (F) of the indicated genes in granulosa cells treated with DMSO or EPZ015666 (5μM) for 5 days. The protein expression in Western blot analysis was quantified and normalized to that of GAPDH (B and E). The data are presented as the mean±SEM. For B,E,F, n=3; For C, n=5. *, P < 0.05. **, P < 0.01.

### The expression of WT1 was regulated by PRMT5 at the translational level

PRMT5 has been reported to regulate the translation of several genes in an IRES-dependent manner^17, 18^. Internal ribosome entry sites (IRESs) are secondary structures in the 5’UTR that directly recruit the ribosome cap independently and initiate translation without cap binding and ribosome scanning^19–21^. *Wt1* 5’UTR is 268bp, GC-rich (68%) and contains 7 CUG codons and 1 AUG codon. These features usually act as strong barriers for ribosome scanning and conventional translation initiation. Translation initiation in a number of these mRNAs is achieved via IRES-mediated mechanisms^21^. To test whether the *Wt1* 5’UTR contains an IRES element, we utilized a pRF dicistronic reporter construct in which upstream Renilla luciferase is translated cap-dependently, whereas downstream firefly luciferase is not translated unless a functional IRES is present. A stable hairpin structure upstream of Renilla luciferase minimizes cap-dependent translation^19^ (Fig. 6A). *Wt1* 5’UTR was inserted into the intercistronic region between Renilla and firefly luciferase (named pRWT1F), and primary granulosa cells were transfected with pRF or pRWT1F. The firefly/Renilla luciferase activity ratio was analyzed 48 hours later. As shown in Fig. 6B, the firefly/Renilla luciferase activity ratio was dramatically increased in pRWT1F-transfected cells compared to pRF-transfected cells. In contrast, the firefly/Renilla luciferase activity ratio was not increased when *Wt1* 5’UTR was inserted in the reverse direction (pRWT1-RevF) (Fig. 6B). The luciferase activity was dramatically increased with insertion of *Ccnd1* 5’UTR as a positive control, which has been reported to contain an IRES element in the 5’UTR^22^. These results suggest that *Wt1* 5’UTR probably contains an IRES element.

**Figure 6.**
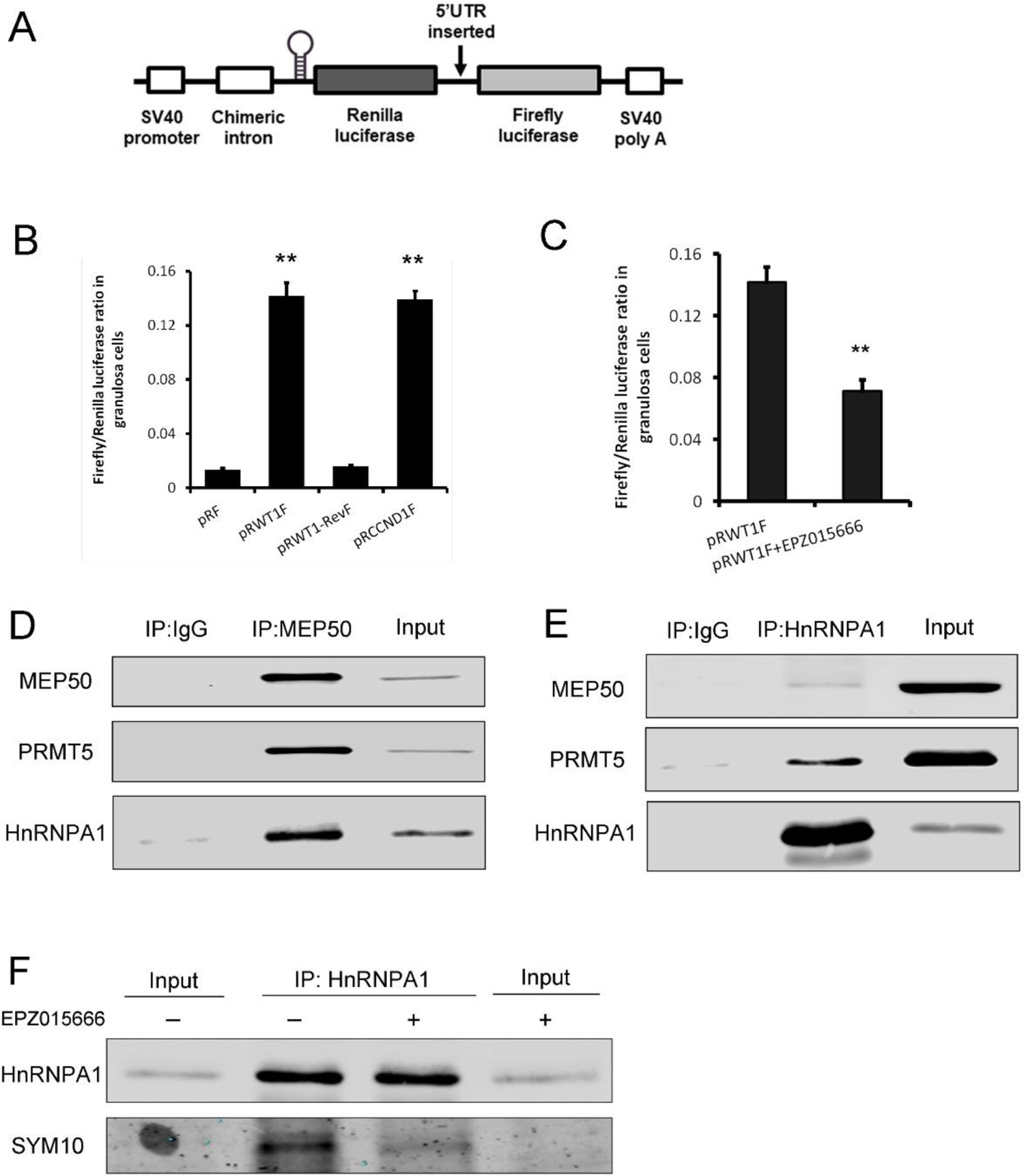
PRMT5 regulated translation of *Wt1* mRNA by inducing IRES activity in the 5’UTR. A, Schematic representation of the dicistronic reporter construct. B, *Wt1* 5’UTR has IRES activity. Cultured primary granulosa cells were transfected with pRF, pRWT1F (pRF with the *Wt1* 5’UTR inserted), pRWT1-RevF (pRF with the *Wt1* 5’UTR inserted in reverse orientation), or pRCCND1F (pRF with the *Ccnd1* 5’UTR inserted). The firefly and Renilla luciferase activities were measured 24 hours later, and the ratios of firefly luciferase activity to Renilla luciferase activity were calculated. C, Luciferase activity was decreased in primary granulosa cells treated with the PRMT5 inhibitor EPZ015666. Isolated granulosa cells were treated with DMSO or EPZ015666 for 4 days. The day granulosa cells were isolated was denoted as day 1. On day 4, granulosa cells were transfected with pRWT1F. Twenty-four hours later, the cells were harvested for luciferase activity analysis. The ratios of firefly luciferase activity to Renilla luciferase activity were calculated. In A-C, the data are presented as the mean±SEM, n=4. **, P < 0.01. D, HnRNPA1 was pulled down with an antibody against the PRMT5-associated protein MEP50. E, PRMT5 and MEP50 were pulled down by an HnRNPA1 antibody. F, The symmetric dimethylation of HnRNPA1 was decreased after EPZ015666 treatment in primary granulosa cells.

To verify that firefly luciferase protein was synthesized by translation of an intact dicistronic transcript instead of a monocistronic mRNA generated by cryptic splicing or promoter within the dicistronic gene^23^, mRNA from pRF- or pRWT1F-transfected cells was treated with DNase, reverse-transcribed, and then amplified with primers binding to the 5’ end of Renilla luciferase and 3’ end of firefly luciferase open reading frame spanning the whole transcript. Only one band was detected in both cells with the expected molecular weight (Fig. S5A). Moreover, qPCR analysis of firefly and Renilla luciferase mRNA levels also showed that the firefly/Renilla luciferase mRNA ratio was not different between pRF- and pRWT1F-transfected cells (Fig. S5B), further excluding the possibility that insertion of the *Wt1* 5’UTR into pRF generated a monocistronic firefly ORF.

**Figure S5.**
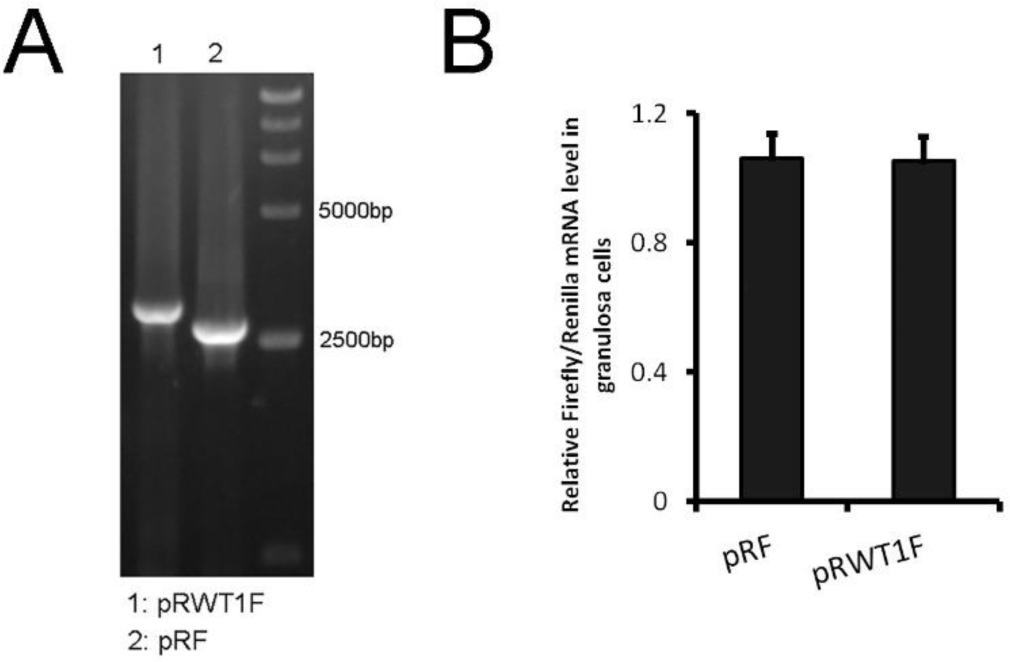
The increased luciferase activity of pRWT1F was not due to a monocistronic firefly ORF generated by cryptic splicing or promoter within the dicistronic gene. A, Primary granulosa cells were transfected with pRF or pRWT1F plasmids. RNA was isolated, DNase-treated, reverse-transcribed, and amplified using PCR primers that bind to the 5’ end of Renilla luciferase and the 3’ end of the firefly luciferase sequence (A) or processed for real-time PCR assays to analyze firefly and Renilla luciferase mRNA levels (B). The expression of firefly luciferase mRNA was normalized to that of Renilla luciferase mRNA. The data are presented as the mean ± SEM, n=3.

To investigate the effect of PRMT5 on *Wt1* IRES activity, granulosa cells were treated with EPZ015666 for 4 days, we found *Wt1* IRES activity was decreased significantly by EPZ015666 (Fig. 6C). These results indicate that PRMT5 regulates *Wt1* expression at the translational level through inducing its IRES activity in granulosa cells.

### Wt1 IRES activity was regulated by PRMT5 through methylation of HnRNPA1

IRES-mediated translation depends on IRES *trans*-acting factors (ITAFs), which function by associating with the IRES and either facilitate the assembly of initiation complexes or alter the structure of the IRES^23, 24^. Heterogeneous nuclear ribonucleoprotein A1 (HnRNPA1) is a well-studied RNA binding protein that plays important roles in pre-mRNA and mRNA metabolism^25^. HnRNPA1 is also an ITAF that has been reported to regulate the IRES-dependent translation of many genes, such as *Ccnd1*, *Apaf1*^26^, *Myc*^24^, *Fgf2*^27^, and *Xiap*^28, 29^. HnRNPA1 can be methylated by PRMT1^28^ or PRMT5^17, 18^, which regulates the ITAF activity of HnRNPA1. To test whether PRMT5 interacts with HnRNPA1 in granulosa cells, coimmunoprecipitation experiments were conducted. We found that HnRNPA1 and PRMT5 were pulled down by antibody against the PRMT5 main binding partner MEP50 (Fig. 6D). Conversely, PRMT5 and MEP50 could be pulled down by the HnRNPA1 antibody (Fig. 6E). We also found that the level of symmetric dimethylation of HnRNPA1 was significantly reduced with EPZ015666 treatment in granulosa cells (Fig. 6F).

To test whether HnRNPA1 functions during PRMT5-mediated *Wt1* translation, HnRNPA1 was knocked down in granulosa cells via siRNA transfection. Western blot analysis results showed that HnRNPA1 protein levels were significantly decreased with siRNA transfection (Fig. 7A). We found that WT1 protein level was increased significantly in granulosa cells after knockdown of HnRNPA1. The decreased WT1 expression in EPZ015666-treated granulosa cells was partially reversed by knockdown of HnRNPA1 (Fig. 7A). The luciferase activity of pRWT1F was increased in granulosa cells with HnRNPA1 siRNA treatment and decreased in those with EPZ015666 treatment. The decreased luciferase activity in EPZ015666-treated granulosa cells was partially reversed by knockdown of HnRNPA1 (Fig. 7B). To further confirm the effect of HnRNPA1 on *Wt1* IRES activity, HnRNPA1 was overexpressed in granulosa cells, and we found that *Wt1* IRES activity was significantly decreased (Fig. 7D). These results indicated that as an ITAF, the effect of HnRNPA1 on *Wt1* IRES activity was repressive.

**Figure 7.**
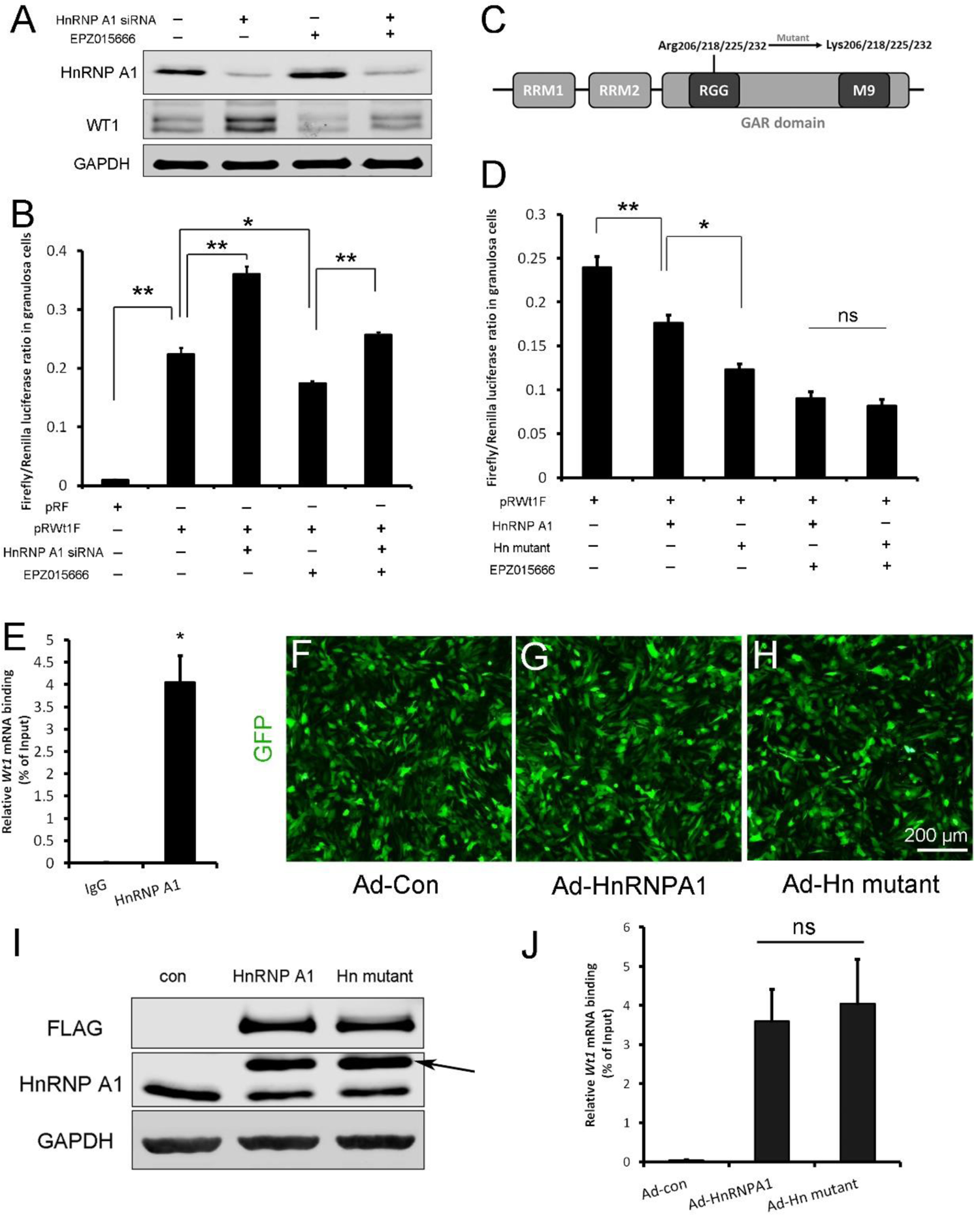
*Wt1* IRES activity is regulated by PRMT5 via methylation of HnRNPA1. A, Western blot analysis of HnRNPA1 and WT1 in granulosa cells after HnRNPA1 siRNA transfection or EPZ015666 treatment. B, Luciferase activity analysis of pRWT1F in granulosa cells after HnRNPA1 siRNA transfection or EPZ015666 treatment. Isolated granulosa cells were treated with DMSO or EPZ015666 for 4 days. The day granulosa cells isolated was denoted as day 1. On day 2, cells were transfected with control siRNA or siRNA to HnRNPA1. 48h later, pRF or pRWT1F were transfected. The luciferase activity of pRWT1F was calculated as the ratio of firefly luciferase activity to Renilla luciferase activity. C, Schematic diagram of HnRNPA1 protein domains. HnRNPA1 contains two RRMs (RNA recognition motifs). The glycine/arginine-rich (GAR) domain contains an RGG (Arg-Gly-Gly) box and a nuclear targeting sequence (M9). Four arginine residues within the RGG motif were mutated to lysine. D, Luciferase activity analysis of pRWT1F in granulosa cells after EPZ015666 treatment or overexpressing HnRNPA1 or arginine-mutated HnRNPA1. Isolated granulosa cells were treated with DMSO or EPZ015666 for 4 days. On day 3, flag-tagged HnRNPA1 or mutant plasmids were cotransfected with pRWT1F into granulosa cells. 48h later, cells were harvested for luciferase activity analysis. E, RNA immunoprecipitation was conducted in granulosa cells using an HnRNPA1 antibody, and the *Wt1* mRNA pulled down by HnRNPA1 was analyzed with real-time PCR. F-H, Primary granulosa cells were cultured and infected with control, flag-tagged HnRNPA1, or mutant HnRNPA1 (Ad-Hn mutant) adenoviruses. The expression of control and mutant HnRNPA1 was examined by Western blot analysis (I). J, RNA immunoprecipitation was conducted using a FLAG antibody, and *Wt1* mRNA pulled down by control or mutant HnRNPA1 protein was analyzed with real-time PCR. For B, D (n=4) and E, J (n=3), the data are presented as the mean±SEM. *, P < 0.05. **, P < 0.01.

There are five arginine residues in the HnRNPA1 glycine/arginine-rich (GAR) motif, which can be symmetrically or asymmetrically dimethylated by PRMT5^17^ or PRMT1^28, 30^, respectively. R206, R218, R225 and R232 are required for HnRNPA1 ITAF activity^17, 28^. To determine the role of HnRNPA1 arginine methylation in *Wt1* IRES activity, the four arginine residues were mutated to lysines (Fig. 7C), and flag-tagged HnRNPA1 or mutant plasmids were cotransfected with pRWT1F into granulosa cells. We found that *Wt1* IRES activity was further decreased in granulosa cells overexpressing mutant HnRNPA1 compared to those overexpressing wild-type HnRNPA1 (Fig. 7D). However, the difference in *Wt1* IRES activity between cells overexpressing mutant HnRNPA1 and cells overexpressing wild-type HnRNPA1 disappeared when the granulosa cells were treated with EPZ015666 (Fig. 7D). These results indicate that the repressive function of HnRNPA1 on *Wt1* IRES activity is inhibited by PRMT5-mediated arginine symmetric dimethylation.

To test the interaction between HnRNPA1 and *Wt1* mRNA, RNA immunoprecipitation was performed with an HnRNPA1 antibody in primary granulosa cells. As shown in Fig. 7E, *Wt1* mRNA was pulled down by the HnRNPA1 antibody in granulosa cells. Next, granulosa cells were infected with flag-tagged wild-type or arginine-mutant HnRNPA1 adenovirus (Fig. 7F-I) and RNA immunoprecipitation was conducted with an FLAG antibody. The results showed that mutation of arginines did not affect the interaction between HnRNPA1 and *Wt1* mRNA (Fig. 7J).

### The upregulation of steroidogenic genes in Prmt5^flox/flox^;Sf1-cre granulosa cells was repressed by Wt1 overexpression

To test whether the upregulation of steroidogenic genes in *Prmt5*-deficient granulosa cells is due to downregulation of WT1, granulosa cells from *Prmt5^flox/flox^;Sf1-cre* mice were infected with control or GFP-tagged WT1-expressing adenovirus (Fig. 8A, B). *Wt1* protein (Fig. 8D arrow, E) and mRNA (Fig. 8C) levels were dramatically increased in *Prmt5*-deficient granulosa cells after *Wt1* overexpression. We found the expression of steroidogenic genes was significantly decreased in these cells. These results suggest that the aberrant differentiation of *Prmt5*-deficient granulosa cells can be rescued by WT1.

**Figure 8.**
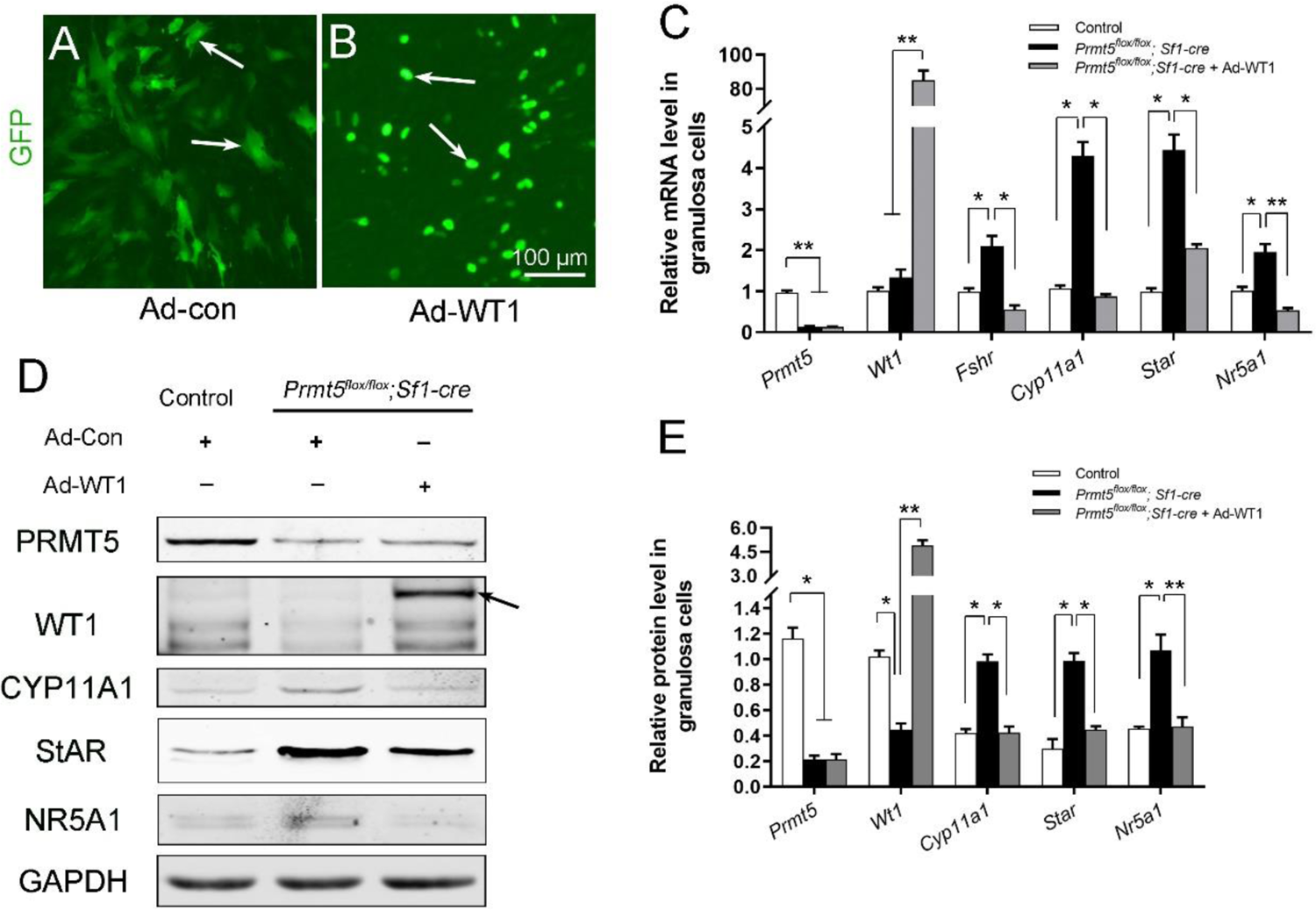
The upregulation of steroidogenic genes in *Prmt5^flox/flox^;Sf1-cre* granulosa cells was reversed by *Wt1* overexpression. A, B, Granulosa cells isolated from control and *Prmt5^flox/flox^;Sf1-cre* mice were cultured and infected with control or GFP-fused *Wt1* adenovirus. The expression of steroidogenic genes was examined by RT-qPCR (C) and Western blot analysis (D). The protein expression in Western blot analysis was quantified and normalized to that of GAPDH (E). C,E, the data are presented as the mean ± SEM (n=3). *, P < 0.05. **, P < 0.01.

## Discussion

Protein arginine methylation is one of the most important epigenetic modifications and is involved in many cellular processes. It has been demonstrated that PRMT5 is required for germ cell survival. In this study, we found that protein arginine methylation plays important roles in granulosa cell development. The development of ovarian follicles is a dynamic process. With follicle development, the morphology and gene expression of granulosa cells are changed. When primordial follicles are activated to grow, the flattened granulosa cells become more epithelial through processes such as cuboidalization and polarization in primary follicles. The granulosa cells in antral follicles express gonadotropin receptors. Before ovulation, granulosa cells begin to express steroidogenic enzymes that are necessary for progesterone and estradiol synthesis^6, 31^. WT1 is expressed at high levels in granulosa cells of primordial, primary, and secondary follicles but decreases with follicle development, suggesting it might be a repressor of ovarian differentiation genes in the granulosa cells^7^. Our previous study demonstrated that the *Wt1* gene is required for lineage specification and maintenance of granulosa cells^8, 9^. Inactivation of *Wt1* causes the transformation of pregranulosa cells to steroidogenic cells. In this study, we found *Prmt5*-deficient granulosa cells began to express steroidogenic genes in secondary follicles and the upregulation of the steroidogenic genes in *Prmt5*-deficient granulosa cells was reversed by *Wt1* overexpression, indicating that PRMT5 is required for preventing the premature differentiation of granulosa cells via regulation of WT1 expression. Coordinated interaction between granulosa cells and oocytes is required for successful follicle development and production of fertilizable oocytes. The premature luteinized granulosa cells will lose their structural and nutritional support for oocytes which will lead to follicle growth arrest or atresia at early stages of folliculogenesis.

Our previous study demonstrated that WT1 represses *Sf1* expression by directly binding to the *Sf1* promoter region and that inactivation of *Wt1* causes upregulation of *Sf1,* which in turn activates the steroidogenic program^8^. In the present study, the mRNA and protein levels of *Sf1* were significantly upregulated after WT1 loss; therefore, the upregulation of steroidogenic genes in *Prmt5*-deficient granulosa cells is due to the increased expression of *Sf1*.

As an important nuclear transcription factor, the function of WT1 in granulosa cell development has been investigated. However, the molecular mechanism that regulates the expression of this gene is unknown. In this study, we found that the expression of WT1 at the protein level was dramatically reduced in *Prmt5*-deficient granulosa cells, whereas the mRNA level was not changed, indicating that PRMT5 regulates *Wt1* expression at the posttranscriptional level. In our mouse model, *Prmt5* was inactivated in granulosa cells at the early embryonic stage. However, defects in follicle development were not observed until 2 weeks after birth. Follicle development was arrested. This outcome probably occurred because *Prmt5* is not expressed in granulosa cells before the development of primary follicles (Fig. S1). During the early stage, *Wt1* expression is also maintained in pre-granulosa cells, therefore, we speculate there must be another factor(s) regulating *Wt1* expression before primary follicle stage.

More than 100 mRNAs in mammals contain IRES elements in their 5’UTRs^32^, which are involved in various physiological processes, such as differentiation, cell cycle progression, apoptosis and stress responses^33^. The 5’UTR sequence of *Wt1* mRNA is highly conserved, with more than 85% homology among the sequences of 29 mammalian species. Our study indicates that the *Wt1* 5’UTR has IRES activity. HnRNPA1 belongs to the HnRNP family, which comprises at least 20 members associated with RNA processing, splicing, transport and metabolism^33, 34^. As a main ITAF, HnRNPA1 either activates the translation of *Fgf2*^27^, *Srebp-1a*^35^, and *Ccnd1*^22^ or inhibits the translation of *Xiap*^29^, *Apaf*^26^, and *Bcl-xl*^36^. The underlying mechanism by which HnRNPA1 activates some IRESs but suppresses other IRESs is still unknown. HnRNPA1 may compete with other ITAFs for binding or may modify IRES structure and thus regulate IRES activity^26, 29^.

It has been reported that the expression of several genes is regulated by PRMT5 at the protein level^17, 37^. *Gao et al.* reported that PRMT5 regulates IRES-dependent translation via methylation of HnRNPA1 in the 293T and MCF-7 cell lines. They found that HnRNPA1 activates the IRES-dependent translation and that methylation of HnRNPA1 facilitates the interaction of HnRNPA1 with IRES mRNA to promote translation^17^. In the present study, we found that *Wt1* IRES activity was repressed by HnRNPA1 (Fig. 9A) and that the repressive effect of HnRNPA1 was reversed by PRMT5-mediated arginine methylation; thus, *Wt1* IRES-dependent translation was promoted by PRMT5 (Fig. 9B).

**Figure 9.**
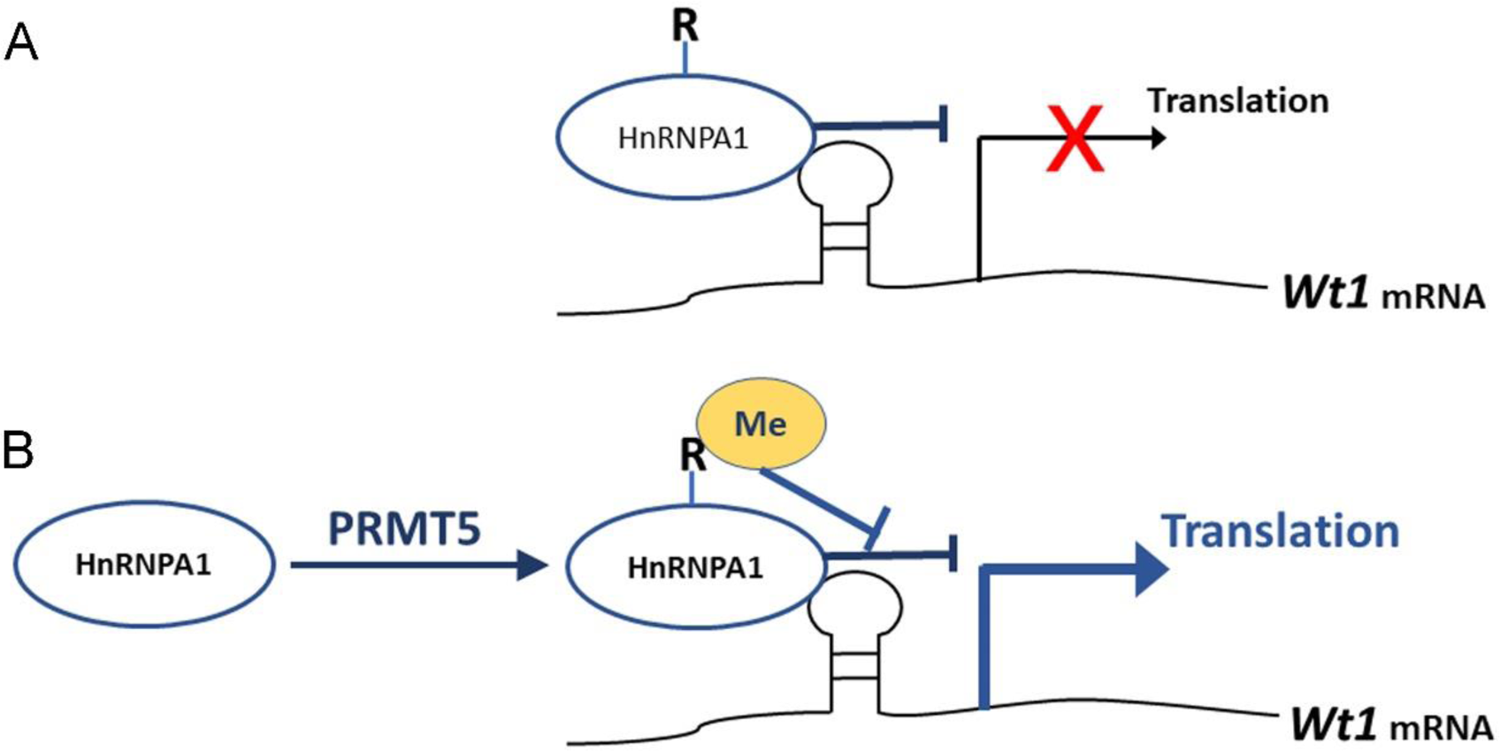
Schematic illustration of how PRMT5 regulates *Wt1* mRNA translation. A, As an ITAF, HnRNPA1 binds to *Wt1* mRNA and inhibits the IRES-dependent translation of *Wt1*. B, PRMT5 catalyzes symmetric methylation of HnRNPA1, which suppresses the ITAF activity of HnRNPA1 and promotes the translation of *Wt1* mRNA. R, arginine. Me, methylation.

The ITAF activity of HnRNPA1 can be regulated by posttranslational modifications^33^. Phosphorylation of HnRNPA1 on serine 199 by Akt inhibits IRES-dependent translation of *c-myc* and *cyclin D1*^22, 24^. Symmetric dimethylation of HnRNPA1 by PRMT5 enhances HnRNPA1 ITAF activity and promotes the translation of target mRNAs^17^. Asymmetric dimethylation of HnRNPA1 by PRMT1 inhibits its ITAF activity^28^. These results suggest that arginine methylation has different effects on the ITAF activity of HnRNPA1 according to different IRESs and cell contexts. Our study demonstrated that HnRNPA1 ITAF activity toward *Wt1* mRNA was repressed by PRMT5-mediated arginine methylation. However, the affinity between HnRNPA1 and *Wt1* mRNA was not affected after mutation of arginine residues, consistent with the findings of a previous study^28^. Therefore, the inhibition of HnRNPA1 ITAF activity by PRMT5 does not occur through changes in the binding of HnRNPA1 to *Wt1* mRNA. The underlying mechanism needs further investigation.

Epigenetic modification is involved in numerous cellular processes. However, the functions of epigenetic modification in granulosa cell development have not been well studied. In this study, we demonstrated that *Prmt5* is required for maintenance of granulosa cell identity in follicle development and that inactivation of *Prmt5* causes premature luteinization of granulosa cells. Our study also demonstrates that PRMT5 regulates WT1 expression at the translational level by methylating HnRNPA1. This study provides very important information for better understanding the regulation of gonad somatic cell differentiation.

## Materials and Methods

### Mice

All animal experiments were carried out in accordance with the protocols approved by the Institutional Animal Care and Use Committee at the Institute of Zoology, Chinese Academy of Sciences (CAS). All mice were maintained on a C57BL/6;129/SvEv mixed background. *Prmt5^flox/flox^; Sf1-cre* female mice were obtained by crossing *Prmt5^flox/flox^* mice with *Prmt5^+/flox^; Sf1-cre* mice. *Prmt5^flox/flox^* and *Prmt5^+/flox^* female mice were used as controls.

### Plasmid and adenovirus

The dicistronic construct pRF was a generous gift from Professor Anne Willis, University of Cambridge. pRWT1F, pRCCND1F and pRWT1-RevF were constructed by inserting the mouse *Wt1* 5’UTR, human *Ccnd1* 5’UTR or mouse *Wt1* 5’UTR in reverse orientation into EcoRI and NcoI sites of the pRF vector. Mouse *Wt1* 5’UTR and human *Ccnd1* 5’UTR sequence were amplified by PCR and the primers used were: pRWT1F-F: CCGGAATTCTGTGTGAATGGAGCGGCCGAGCAT, pRWT1F-R: CTAGCCATGGGATCGCGGCGAGGAGGCG; pRWT1-RevF-F: CTAGCCATGGTGTGTGAATGGAGCGGCCGAGCAT, pRWT1-RevF-R: CCGGAATTCGATCGCGGCGAGGAGGCG; pRCCND1F-F: GCTGAATTCCACACGGACTACAGGGGAGTTTT, pRCCND1F-R: CGGCCATGGGGCTGGGGCTCTTCCTGGGC. The primers amplifying the whole transcript of pRF binding to the 5’ end of renilla and 3’ end of firefly ORF: pRF-F: GCCACCATGACTTCGAAAGTTTATGA; pRF-R: TTACACGGCGATCTTTCCGC.

FLAG-tagged HnRNPA1 and mutant plasmids were generated by inserting the coding sequence and a mutant sequence of mouse HnRNPA1, respectively, into NheI and BamHI sites of the pDC316-mCMV-ZsGreen-C-FLAG vector. HnRNPA1-F: CTAGCTAGCCACCATGTCTAAGTCCGAGTCTCCCAAGGA, HnRNPA1-R: CGCGGATCCGAACCTCCTGCCACTGCCATAGCTA. Adenoviruses containing WT1 coding sequence, HnRNPA1 or the mutant sequence were generated using the Gateway Expression System (Invitrogen).

### Isolation of granulosa cells, transient transfection, infection and luciferase assay

Granulosa cells were isolated from mice at 16-18 days old. After mechanical dissection, ovaries were cut into several parts and incubated in PBS containing 1 mg/ml collagenase IV (Sigma) in a water bath with circular agitation (85 rpm) for 5 min at 37°C. Follicles were allowed to settle and washed in PBS. The supernatant were discarded. A secong enzyme digestion was performed in PBS containing 1 mg/ml collagenase IV, 1 mg/ml Hyaluronidase, 0.25% Trypsin, and 1mg/ml DNase I (Applichem) for 15min. FBS was added to stop the digestion and cell suspension was filtered through a 40-μm filter. Cells were centrifuged, washed and then plated in 24-well plate in DMEM/F12 supplemented with 5% FBS. For EPZ015666 treatment, granulosa cells were incubated in the medium with the addition of 5μM EPZ015666 (MedChemExpress) for 4-5 days. When cells were approximately 70% confluent, granulosa cells were transfected with plasmids or infected with adenovirus according to the experiments. At the end of culture, cells were lysed for RT-qPCR, Western blot analysis, or luciferase activity analysis using a dual luciferase reporter assay system (Promega).

Control siRNA or siRNA to HnRNPA1 was purchased from ThermoFisher (S67643, S67644) and transfected into granulosa cells with Lipofectamine 3000 transfection reagent without P3000. 48 hours later, pRF or pRWT1F were transfected and luciferase activities were measured the following day.

### In vitro ovarian follicle culture

Follicles were dissected and cultured as previously described^38^. Briefly, ovaries of 14-day-old mice were dissected aseptically using the beveled edges of two syringe needles. Follicles with 2–3 layers of granulosa cells, a centrally placed oocyte, an intact basal membrane and attached theca cells were selected and cultured individually in 20 μl droplets of culture medium (αMEM supplemented with 5% FBS, 1% ITS and 100 mIU/ml recombinant FSH). The culture were maintained in 37℃ and 5% CO2 in air. The medium was replaced every other day. The morphology of the follicles was recorded daily under a microscope.

### Co-immunoprecipitation

Granulosa cells isolated from mice at 16-18 days old were cultured in 10-cm dishes and lysed with lysis buffer (50mM Tris·HCl (pH 7.5), 150 mM NaCl, 1mM EDTA, 1% Nonidet P-40) supplemented with protease inhibitors cocktail (Roche) and 1mM PMSF. One milligram of protein were first pre-cleared with protein G agarose beads (GE) for 1 hour at 4℃, then incubated with 1.5μg of IgG (mouse, Santa Cruz, sc-2025; rabbit, Abmart, B30011S), HnRNPA1 antibody (Abcam, ab5832), or MEP50 antibody (Abcam, ab154190) for 4 hours at 4℃. Then protein A and G agarose beads were added and incubated overnight. The immunoprecipitates were washed 4 times in lysis buffer supplemented with cocktail and PMSF, resolved in loading buffer, incubated for 5 min at 95℃, and then analyzed by Western blotting. The antibodies used in Western blotting include: PRMT5 (Millipore, 07-405), SYM10 (Millipore, 07-412), HnRNPA1 (Abcam, ab5832), MEP50 (Abcam, ab154190).

### Western blot analysis

Granulosa cells were washed with PBS, lysed with RIPA buffer (50 mM Tris– HCl (pH 7.5), 150 mM NaCl, 1% NP-40, 0.1% SDS, 1% sodium deoxycholate, 5mM EDTA) supplemented with protease inhibitors cocktail (Roche) and 1mM PMSF. 30μg total protein was separated by SDS/PAGE gels, transferred to nitrocellulose membrane, probed with the primary antibodies. The images were captured with the ODYSSEY Sa Infrared Imaging System (LI-COR Biosciences, Lincoln, NE, USA). The antibodies used were: PRMT5 (Millipore, 07-405), MEP50 (Abcam, ab154190), WT1 (Abcam, ab89901), FOXL2 (Abcam, ab5096), CYP11A1 (Proteintech, 13363-1-AP), StAR (Santa Cruz, sc-25806), SF1 (Proteintech, 18658-1-AP), FLAG (Sigma, F1804).

### RNA immunoprecipitation

Granulosa cells were isolated from mice at 16-18 days old and cultured in 10-cm dishes. The cells were then lysed with RIP buffer (50 mM Tris-HCl pH7.5, 150 mM NaCl, 5mM EDTA, 1% NP-40, 0.5% sodium deoxycholate) supplemented with protease inhibitor cocktail and 200U/ml RNase inhibitor. 5% of the cell lysate supernatants were used as the input and the remainings were incubated with 1.5 μg of IgG (mouse, Santa Cruz, sc-2025), HnRNPA1 antibody (Abcam, ab5832), or FLAG antibody (Sigma, F1804) for 4 hours at 4℃. Then protein A and G agarose beads were added to immunoprecipitate the RNA/protein complex. The conjugated beads were thoroughly washed with lysis buffer (50 mM Tris–HCl pH7.5, 500 mM NaCl, 5mM EDTA, 1% NP-40, 0.5% sodium deoxycholate) supplemented with cocktail and 200U/ml RNase inhibitor. Bound RNA was extracted using a RNeasy Kit and analyzed with RT-qPCR analysis.

### Real-time RT-PCR

Total RNA was extracted using a RNeasy Kit (Aidlab, RN28) in accordance with the manufacturer’s instructions. One micrograms of total RNA was used to synthesize first-strand cDNA (Abm, G592). cDNAs were diluted and used for the template for real-time SYBR Green assay. *Gapdh* was used as an endogenous control. All gene expression was quantified relative to *Gapdh* expression. The relative concentration of the candidate gene expression was calculated using the formula 2^-ΔΔCT^. Primers used for the RT-PCR are listed in Table S1.

### Immunohistochemistry and immunofluorescence analysis

Immunohistochemistry procedures were performed as described previously^39^. Stained sections were examined with a Nikon microscope, and images were captured by a Nikon DS-Ri1 CCD camera. For immunofluorescence analysis, the 5-μm sections were incubated with 5% BSA in 0.3% Triton X-100 for 1 hours after rehydration and antigen retrieval. The sections were then incubated with the primary antibodies for 1.5 hours and the corresponding FITC-conjugated donkey anti-goat IgG (1:150, Jackson ImmunoResearch) and Cy^TM^3-conjugated donkey anti-rabbit IgG (1:300, Jackson ImmunoResearch) for 1 hour at room temperature. The following primary antibodies were used: WT1 (Abcam, ab89901), FOXL2 (Abcam, ab5096), CYP11A1 (Proteintech, 13363-1-AP), SF1 (Proteintech, 18658-1-AP), 3β-HSD (Santa Cruz Biotechnology, sc-30820). After being washed three times in PBS, the nuclei were stained with DAPI. The sections were examined with a confocal laser scanning microscope (Carl Zeiss Inc., Thornwood, NY).

### Statistical analysis

All experiments were repeated at least three times. For immunostaining, one representative picture of similar results from three to five control or *Prmt5*-deficient ovaries at each time point is presented. The quantitative results are presented as the mean±SEM. Statistical analyses were conducted using GraphPad Prism version 9.0.0. Unpaired two-tailed Student’s t-tests were used for comparison between two groups. For three or more groups, data were analyzed using one-way ANOVA. P-values < 0.05 were considered to indicate significance.

## Data availability

Source data for Fig.5, 6, 7, 8 and Fig. S5 have been provided as Supplementary file 1.

## Acknowledgments

We thank Prof. Anne Willis (University of Cambridge) for her generous gift of the dicistronic contruct pRF. We thank Prof. Humphrey Hung-Chang Yao (NIEHS/NIH) for the *Sf1-cre* mice. This work was supported by National key R&D program of China (2018YFC1004200, 2018YFA0107700); Strategic Priority Research Program of the Chinese Academy of Sciences (XDB19000000); The National Science Fund for Distinguished Young Scholars (81525011); The National Natural Science Foundation of China (31970785, 31601193, and 31671496).

## Author contributions

F.G. and M.C. designed the experiments and wrote the manuscript. M.C. and F.F.D. performed the experiments and analyzed the data. All authors discussed the results and edited the manuscript. The authors declare that they have no conflict of interest.

---

**Table S1.**
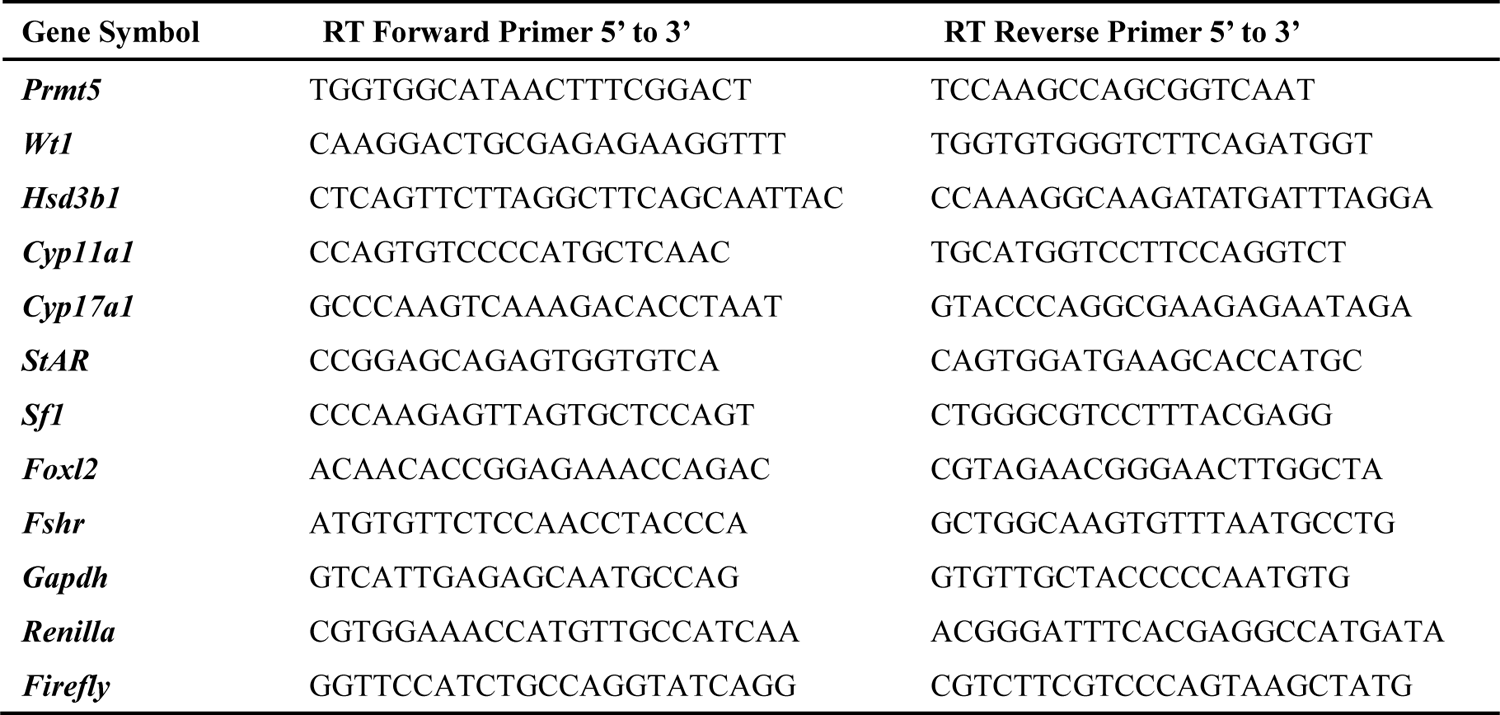
Primers used for real-time PCR analysis.

